# Social evolution and absence of olfactory function in larval honey bees

**DOI:** 10.1101/2025.03.31.646396

**Authors:** Tianfei Peng, Zhenqing Chen, Amy C. Cash Ahmed, Qianlu Feng, Seokjin Yeo, Hee-Sun Han, Gene E. Robinson

## Abstract

Social evolution made larval honey bees dependent on adult colony members for feeding. We therefore predicted they have diminished olfactory capabilities, and based on organismal resource conservation theory, also have downregulated olfactory receptor (OR) gene expression. Behavioral assays demonstrated that larvae cannot find food via olfaction and expressed very low levels of *Orco*, an essential gene for OR function. By contrast, larvae showed higher expression of *Ir25a,* an essential gene for multiple forms of sensory perception including gustation. These results suggest larvae rely on taste for feeding. In addition, considering that adult bees use OR-based olfaction extensively, they demonstrate strong developmental regulation of the OR system. Comparative transcriptomics of social and non-social insects further highlight the role of social evolution in shaping this sensory trait.

## Main Text

Social evolution has given rise to traits that have had huge impacts on life on this planet, including agriculture, language, manufacturing, construction, and warfare. All of these result from systems of division of labor among society members, and they are seen in both human and insect societies. Social evolution also results in the loss of certain traits among some society members, such as independent food searching behavior by honey bee larvae.

A hallmark of the division of labor in insect societies is cooperative brood care, and in honey bees, this has resulted in larvae that are confined and individually reared in cells in the waxen honeycombs. They do not engage in independent food searching behavior, which by contrast is a common trait of larvae of non-social insect species. Honey bee larvae are visited and fed extensively by alloparenting adult colony members (*1*). Larvae produce pheromones that influence adult behavior, but there is no experimental evidence that larvae perceive chemical information that influences their own development or behavior (*2, 3*). By contrast, adult worker bees rely heavily on chemical communication for food finding and chemical communication; they use canonical olfactory receptors (ORs) and the honey bee genome has a relatively large OR gene repertoire (*4*).

The sensory capabilities of a species are influenced by both the environments it has evolved in and the limitations imposed by developmental and other evolutionary factors. For example, species that live in dark environments, such as cave-dwelling fish, have permanently lost the neural and molecular machinery needed to process visual information, a phenomenon known as regressive evolution (*5*). Because social evolution has rendered larval honey bees dependent upon their older sisters to be fed, we predicted that they lack olfactory capabilities. We also tested the prediction that larvae have a downregulated OR system. This prediction is based on organismal resource conservation theory, extended to the molecular level, which postulates that molecular machinery underlying systems not used during a particular life stage will be downregulated (*6, 7*). Regressive evolution would not be possible in this case because adult bees use OR-based olfaction extensively.

We used behavioral analysis to explore the hypothesis that honey bee larvae cannot orient to food via olfaction. Larvae generally show very little mobility when reared naturally each in their cell. However, when removed from their cell and starved, they move slowly for hours with sideways turns, which results in circular or spiral food searching paths (*1*). Using this information, we modified a laboratory assay developed for *Drosophila melanogaster* larvae (*8*) to use on honey bee larvae. Droplets of standard honey bee larval diet were placed 0.5 cm apart, either in volumes of 2 or 5 µl. The 5 µl droplets were placed closer to each other as well as closer to the larvae, so a higher proportion of volatiles presumably was released (Fig. 1A). If honey bee larvae can smell, they would be predicted to engage more with the 5 µl, rather than 2 µl, droplets, as observed in larvae from other species known to be able to smell (*9, 10*). Extensive observation (5542 hours of observation: 326 individuals, 17 h per individual) revealed there were no differences in the number of 2µl or 5µl food droplets contacted and consumed, measured either cumulatively over time or in total (Totals: LME, F = 0.58653, p = 0.4443) (Fig. 1B and 1C, table S1). In addition, there was no significant correlation between the number of food droplets consumed by a larva and its weight, regardless of food droplet volume (Spearman’s correlation, 2ul: ρ = 0.021, p =0.84; 5ul: ρ = -0.074, p = 0.48) (Fig. 1D, E). The lack of directed movement by honey bee larvae (video S1) is in contrast to the food-directed movement of *Drosophila* larvae (*11*). *Drosophila* larvae receive no care from parents or other adults and must find food themselves. These results indicate that honey bee larvae do not use olfaction to detect food.

**Fig. 1.**
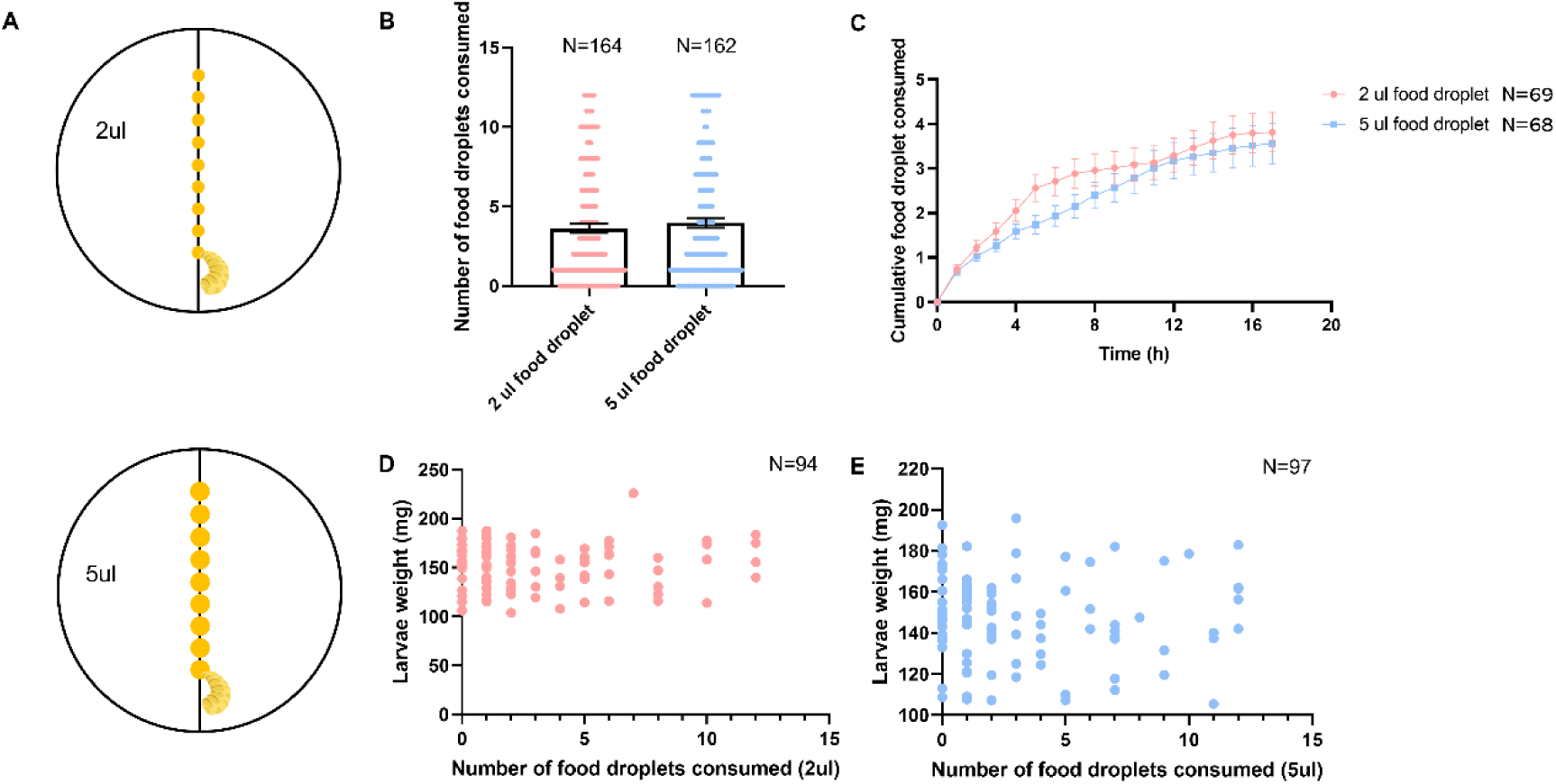
Results of larval honey bee food searching assay. (A) Design of the assay. (B) Total number of food droplets consumed by each larva over 17 h, sample sizes on the graph. Data are shown as mean ± s.e. (C) Cumulative number of food droplets consumed by each larva over 17 h. Detailed statistical analysis is found in Table S3. Data are shown as mean ± s.e. (D) and (E) Relationship between individual initial larval weight and the final number of food droplets it consumed. Each colored dot represents an individual larva.

To determine the mechanism underlying the behavioral results we performed gene expression analysis, focusing on the *OR co-receptor* (*Orco*) and *ionotropic co-receptor 25a* (*Ir25a*) genes. We hypothesized that OR-based gene expression is downregulated in honey bee larvae. Co-receptors for the OR and IR gene families are central to OR and IR function, respectively, with insects possessing only one OR co-receptor (*Orco*) and three IR co-receptors (*Ir8a*, *Ir76b*, and *Ir25a*) conserved across species (*12*). Due to a receptor complex’s reliance on its co-receptor, a mutation in a single co-receptor gene can compromise an insect’ s ability to detect entire classes of environmental sensory cues. Supporting this, CRISPR-induced mutations in *Orco* severely reduced olfactory function and led to asocial behavior in ants (*13, 14*). CRISPR-induced mutations in *Orco* also disrupted the development of the antennal lobes in honey bees (*15*). Therefore, we chose to measure *Orco* expression as an efficient way to assess general OR-based olfactory capabilities, and *Ir25a* expression to assess other forms of food-related sensory perception.

We conducted a comparative analysis of *Orco* and *Ir25a* expression using quantitative polymerase chain reaction (qPCR) in all three honey bee castes: workers, queens, and drones, at three different developmental stages, larval, pupal, and adult. Measurements were made in individual antennae (pupae and adults) or heads (larvae, which have only rudimentary antennae).

*Orco* and *Ir25a* exhibited consistent expression differences across the three castes (Fig. 2, table S2). *Orco* expression was barely detectable in larvae, increasing in pupae, and highest in adults. *Ir25a* also exhibited the lowest expression in larvae, but its expression was significantly higher than *Orco* in larvae. These results are consistent with previously published RNA sequencing results, generated for other purposes (table S3) (*16-22*). Those studies showed low expression levels of all OR genes including *Orco*, while only the IR co-receptor (*Ir25a*) was highly expressed.

**Fig. 2.**
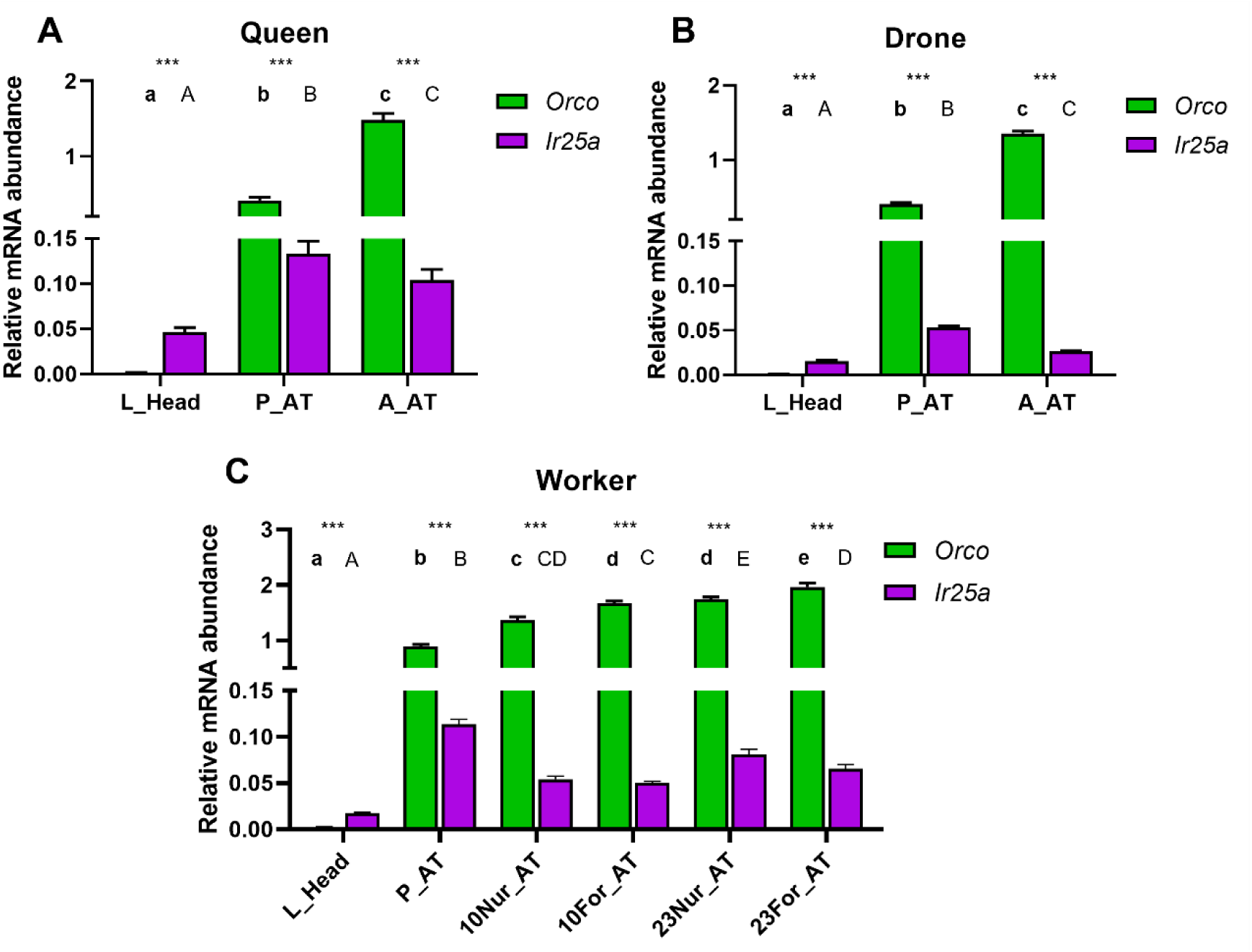
*Orco* and *Ir25a* expression as a function of honey bee caste, development, and behavioral maturation. L_Head = larval head; P_AT = pupal antennae; A_AT = adult antennae (either queen or drone); 10Nur_AT = 10-day-old nurses; 10For_AT = 10-day-old precocious foragers, 23Nur_AT = 23-day-old overage nurses; 23For_AT = 23-day-old foragers. Colonies that produced same-age cohorts of young and old nurse and forager bees were created using a well-established social manipulation (*40*) to differentiate the effects of age and behavior. Lowercase letters denote statistical significance for *Orco* and uppercase letters for *Ir25a*; identical letters denote no significant differences, and different letters denote significant differences. *** denotes significant differences in the expression levels of *Orco* and *Ir25a* within the same tissue (p < 0.0001). Detailed statistical analysis is found in Table S2. Data are shown as mean ± s.e.

Our results also demonstrated that *Orco* expression in adults was dynamically regulated with age and behavioral occupation, consistent with the role of olfaction in the various roles they play that are related to colony growth and development (Fig. 2C, table S2). Adult honey bees use ORs extensively for olfaction (*23*).

It is possible that *Orco* expression occurs in a small subset of cells in the (rudimentary) antennae of larva, and hence the total amount of transcripts is too low to be detected by qPCR. To investigate this possibility, hybridization chain reaction (HCR) RNA fluorescence *in situ* hybridization (RNA-FISH) was performed on sectioned second instar larva (Fig. 3A). Consistent with the qPCR results, the expression of *Orco* was undetectable (Fig. 3B, fig. S1), while robust expression was detected in pupal antenna (fig. S2). By contrast, *Ir25a* expression was readily detected in both larval and pupal tissue (Fig. 3B, fig. S2). These results suggest that honey bee larvae use IRs, but not ORs, to perceive chemical cues.

**Fig. 3.**
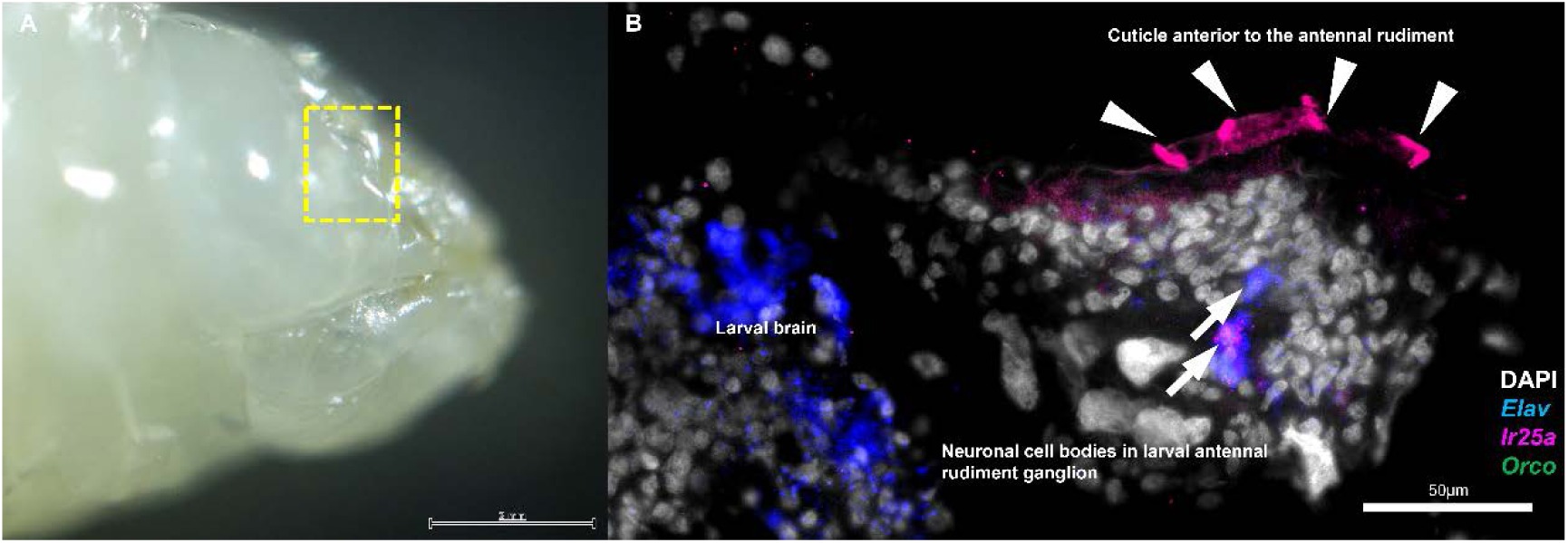
Whole mount HCR RNA-FISH of *Orco* and *Ir25a* expression in the head of honey bee larva. (A) Lateral view of the larval head. The yellow square indicates the antennal rudiment of the larva. (B) Higher magnification of the area boxed in A shows gene expression in a longitudinal section of the larval antennal rudiment. Different colors represent different genes as per legend. *Elav* served as a neuronal marker, and DAPI was used as a stain for nuclear quantitation. Arrows indicate the region of neuronal cell bodies in the larval antennal rudiment ganglion. Arrowheads show the area of the cuticle anterior to the antennal rudiment. No *Orco* signal was detected above background level.

Because ORs are involved exclusively in olfaction whereas IRs are involved in gustation (*24*), these gene expression results support our behavioral findings showing that honey bee larvae can taste but not smell. It is possible that honey bee larvae also use gustatory receptors (GRs); a small number of GRs are present in the honey bee genome (*25*), but their expression levels in larvae are extremely low and the roles of co-receptors in the GR gene family remain unknown (table S3). This means that surveying their involvement in honey bee feeding behavior would not be as efficient as the strategy we used for ORs and IRs, so we did not.

Ecology plays a crucial role in shaping both sensory and social traits in organisms. For example, species that have adopted a parasitic lifestyle exhibit diminished sensory capabilities and a reduction in essential social traits (*26*). By contrast, honey bee larvae show reduced sensory capabilities, apparently because honey bee social evolution has enhanced adult social capacities, including the care and feeding of the larvae.

We also obtained additional evidence for the role of social evolution in shaping larval olfaction by conducting a comparative transcriptomic analysis of *Orco* and *Ir25a* expression in 11 insect species, ranging from eusocial to solitary, based on previously published findings (*16-22, 27-36*). Larvae in three eusocial species (*Monomorium pharaonic*, *Bombus terrestris*, *A. mellifera*), which all exhibit extensive care of larvae by adult colony members, have significantly lower relative ratios of *Orco* to *Ir25a* expression compared to facultatively eusocial (*Megalopta genalis*) and subsocial species (*Ceratina calcarata*), which exhibit less care of larvae by adults. *D. melanogaster*, with no parental care, has a higher relative ratio of *Orco* to *Ir25a* expression (Fig. 4, table S4). In addition, it has previously been reported that knocking out the expression of *Orco* in the embryo stage results in abnormal antennal lobe development in social insects, including honey bees, but not in non-social insects (*13-15, 37*). These qualitative results suggest that the developmental regulation of olfactory gene expression reported here is related to social evolution.

**Fig. 4.**
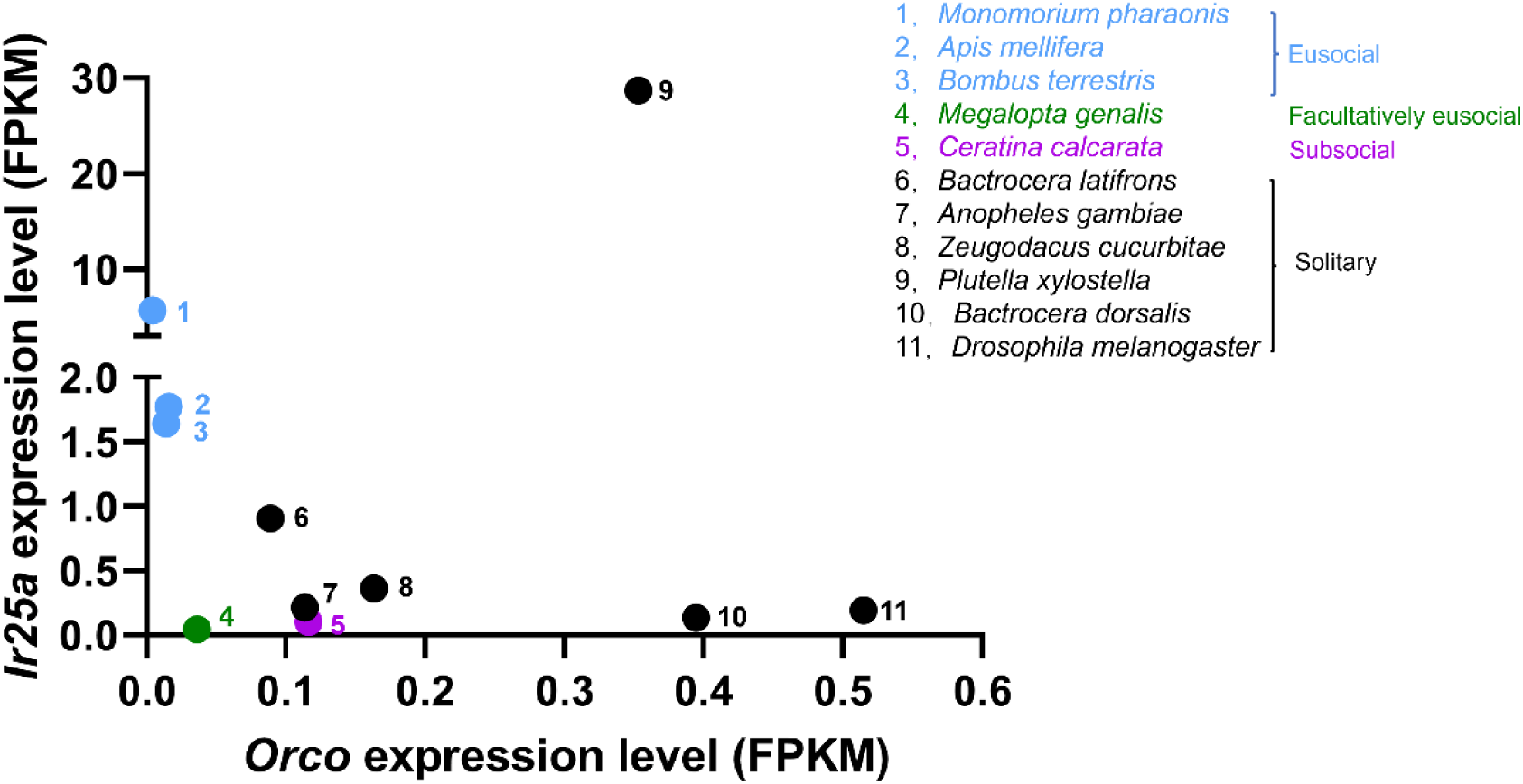
*Orco:Ir25a* expression ratios for larvae from 11 species of insects. For each species, transcriptomic data were gleaned from public databases (from either individual samples or pools of individuals). Sample sizes: *M. pharaonic* = 30; *A. mellifera* = 48; *B. terrestris* = 7; *M. genalis* = 8; *C. calcarata* = 80; *B. latifrons* = 1; *A. gambiae* = 8; *Z. cucurbitae* = 25; *P. xylostella* = 3; *B. dorsalis* = 3; *D. melanogaster* = 198. Different colors represent different types of sociality, from solitary to eusocial. The numbers on the graph adjacent to the data points correspond to the different species, with species names corresponding to the numbers on the right side of the figure. Data sources are given in Supplementary Materials.

Our results are consistent with organismal resource conservation theory and highlight social evolution’s role in shaping the regulation of olfaction. High sensory sensitivity incurs metabolic costs and is expected to occur only when the resulting behavioral response improves fitness (*38, 39*). Because of cooperative brood rearing, honey bee larvae do not need to rely on olfaction for feeding, and this is associated with developmental regulation of key molecular components needed for olfaction. Unlike in other species where evolutionary loss of a sensory ability is associated with the loss or reduction in underlying genes, the present case does not involve gene loss but rather a conservation of molecular resources by developmental reduction in gene expression.

## Acknowledgments

We thank Nathan Beach and Rachel Manweiler for beekeeping assistance, Hongfei Xu and Joseph P. Colgan for useful discussions regarding bumble bee data, and the Robinson lab for valuable comments and feedback on the manuscript. Work involving commercial royal jelly was performed under USDA APHIS PPQ 526 permit # P526P-21-07196.

## Funding

Illinois Sociogenomics Initiative (GER)

National Institutes of Health R35GM147420 (HSH)

## Author contributions

Conceptualization: TFP, ZQC, ACA, QLF, SY, HSH,

GER Methodology: TFP, ZQC, ACA, QLF, SY

Investigation: TFP, ZQC, ACA

Visualization: TFP, ZQC, QLF, SY

Funding acquisition: GER

Project administration: ACA, GER

Supervision: ACA, HSH, GER

Writing – original draft: TFP, GER

Writing – review & editing: TFP, ZQC, ACA, QLF, SY, HSH, GER

## Data availability

All the data can be found in the main text and the supplementary materials. All code and raw data are currently available at https://github.com/nlaptf/honeybee-larva.git.

## Supplementary Materials

**Other Supplementary Materials for this manuscript include the following:**

Table S1 to S5

Table S1: *p*- and *W*-values determined by Wilcoxon Tests

Table S2: *p*-, *F*- and *z*-values determined by linear mixed models

Table S3: The expression level of chemosensory genes in the honey bee larvae

Table S4: *Orco* and *Ir25a* expression levels in different larval species

Table S5: Real-time PCR primers and nucleotides used in experiments

### Materials and Methods

#### Experimental subjects

We used two different types of honey bee colonies as sources of bees for these experiments. For the qPCR analyses, we used individuals from colonies (N = 3) each derived from a queen instrumentally inseminated with semen from a single drone (SDI). For the HCR RNA-FISH (described below), we also used individuals from one of these SDI colonies. For the larval behavior assay, we used individuals from colonies (N = 3) each derived from a naturally mated (NM) queen. All colonies were kept according to standard beekeeping practices at the University of Illinois Bee Research Facility, Urbana, IL, USA. Colonies in this area are a mixture of European subspecies of *Apis mellifera*, primarily *A. mellifera ligustica*.

#### Sample collection

Each colony’s queen was confined to a single empty honeycomb within the hive for 24 h to lay diploid (female worker and queen) and haploid (male “drone”) eggs. These honeycombs were repositioned in their hive to allow the eggs to hatch and larvae to be reared by adult workers, but without access to them by the queen. Four- and five-day-old worker and drone larvae were sampled. For queen larvae, first instar diploid larvae (24 h post-egg hatch) were transferred to commercial plastic “queen-rearing cups” (Mann Lake, USA). They were then collected 72-h post-transfer. Two days before adult eclosion, we removed worker, queen, and drone pupae from their colonies and maintained them in a dark incubator at 34°C, 75% relative humidity. We also collected samples of these penultimate instar pupae for analysis.

The collection process was slightly different for adult drones, queens, and workers sampled for qPCR. Newly eclosed drones were marked on the thorax with enamel paint to indicate their emergence day and source colony. They were then returned to their respective natal colonies and collected after 7 days. Newly eclosed queens were each placed in separate cages to prevent them from showing aggression to each other. These caged queens were subsequently returned to queenless colonies and collected 7 days later. Newly eclosed workers were treated as follows. In order to collect 10 day-old precocious foragers and 23 day-old overage nurses, three single-cohort colonies (SCC) were established (*40*). In brief, we continuously collected newly eclosed workers over a 1- to 3-day period until we had a total of approximately 1200 bees for each SCC. Each bee was then marked on the thorax with enamel paint as an indication of age. Bees were kept in a small hive that contained one honeycomb frame of honey and pollen, one frame of empty comb, one frame of eggs sourced from another colony, and a caged, unrelated queen. The colony was maintained in an incubator until the marking of bees for each colony was completed. The queen was released from her cage, and the colony entrance was opened one day after the colony was placed outside. Collections began when the bees were 7 days old, as precocious foraging behavior typically occurs in bees aged 7 to 10 days (*40*). Over 4 days, we collected precocious foragers; normal-age nurse bees matched in age were collected from inside each colony. When the bees reached the age of 21 days, we similarly collected overage nurses and normal-age foragers also over a period of 4 days. Nurses were identified as they repeatedly inserted their heads into honeycomb cells containing larvae, and forager bees were identified by temporarily blocking the hive entrance and watching for returning bees carrying pollen loads or with a distended abdomen, a mark of nectar foraging (*41*). Larvae, pupae, and adults were flash frozen in liquid nitrogen and then stored at −80°C.

#### Larvae behavior assay

Third instar larvae were removed from three different colonies (each derived from a naturally mated queen). They were removed with a pair of sterile soft tweezers by breaking the wall of their cell on the honeycomb, and gently removing the larvae from the cell. To acclimate the larvae to the odor and taste of Diet C [a standard laboratory diet for larval honey bees: (50% royal jelly, 9% glucose, 9% fructose, 2% yeast extract, and 30% water)] (*15*), they were maintained overnight with this food. Larvae were then placed in plastic cups (1 cm × 1 cm) containing 150 µl of Diet C and incubated for 24 h in a dark incubator at 34°C with 75% relative humidity. Only larvae that remained viable and actively consumed their food were selected for subsequent behavioral assays. Diet C contains a significant amount of royal jelly, the volatiles of which have been shown to elicit electrophysiological responses in nurse bee antennae (*42*). The larva did not show any aversion to diet C and actively fed on it (video S2).

On testing days, we set up multiple Petri dishes (100 mm × 15 mm), each with a 1% agarose-coated lid to maintain humidity, in preparation for the behavioral assays. In the center of the Petri dishes, along the diameter line, we dispensed 12 droplets of a standard formulation of Diet C, with each food droplet center spaced 0.5 cm apart. We divided the Petri dishes into two groups: one group had 5 µl Diet C food droplets, while the other group had 2 µl droplets. Larvae were subjected to 2 h of starvation before being weighed. They then were positioned next to the first food droplet in a Petri dish. All Petri dishes containing larvae were then placed in a dark incubator, maintained at a temperature of 34°C and 75% relative humidity, for 17 h (honey bee larvae move very slowly). A subset of the Petri dishes containing larvae was selected for automated movement tracking. This was done using a Raspberry Pi camera, positioned 30 cm in front of the Petri dishes, under LED lighting. The system was programmed to capture images at 5-min intervals over 17 h (video S1). We then quantified the total and cumulative number of food droplets consumed by each larva as a function of droplet volume.

#### qPCR

We measured the expression of two genes that encode proteins essential to olfaction and gustation in insects, *odorant co-receptor* (*Orco)* and *ionotropic co-receptor (Ir25a)*. *Orco* encodes a highly conserved olfactory co-receptor, which mediates the action of all Ors (*43*); *Ir25a* functions similarly to *Orco*, acting as a co-receptor with various IRs (*44*).

We focused our gene expression analysis on the principal olfactory organs, the antennae of pupae (P_AT) and adults (A_AT); larvae have only rudimentary antennae so we used whole heads (L_Head). Antennae and heads were carefully removed from the bodies, and RNA was extracted using the RNeasy Mini Extraction Kit™ (Qiagen, Germany) according to the manufacturer’s protocol with DNase treatment (Invitrogen, Carlsbad, CA, USA) as has been previously described (*45*), and quantified via a NanoDrop™ spectrophotometer (Thermo Fisher Scientific, USA) and Qubit fluorimeter (Thermo Fisher Scientific, USA). qPCR followed established protocols (*45*). The M-MuLV (New England Biolabs, UAS) reverse transcriptase was used to synthesize the cDNA, Root Cap Protein 1 (*Rcp1*) from *Arabidopsis thaliana* as an external spike-in control to assess synthesis efficiency. qPCR was conducted with SYBR Green dye on an ABI QuantStudio 6 (Thermo Fisher Scientific, USA). We tested seven internal reference genes to find a suitable set for normalization across all categories of bees sampled: *actin 2*, *porphobilinogen deaminase* (*hmbs*), *ribosomal protein 49* (*rp49*), *glyceraldehyde-3-phosphate dehydrogenase* (*gapdh*), *peptidyl-prolyl cis-trans isomerase-like 2* (*ppil2*), *ras-related protein Rab-1A* (*rab1a*), *ribosomal protein L13a* (*rpL13a*). Due to variation in gene expression among larvae, pupae, and adults at different ages, we normalized the candidate genes using different subsets of reference genes for each type of category. For the queen samples, we used the geometric mean of *actin 2*, *gapdh*, and *ppil2* (LME, developmental stages p = 0.25; colony p = 0.87). For the drone samples, we used the geometric mean of *gapdh*, *rpL13a*, and *rab1a* (LME, developmental stages p = 0.10; colony p = 0.08). For the worker samples, we used the geometric mean of *gapdh*, *ppil2*, and *rpL13a* (LME, developmental stages p = 0.68; colony p = 0.40).

We also performed qPCR using second-instar larvae from the same colony used for HCR RNA-FISH (described below) to obtain additional data. The qPCR protocol was the same as described above. Expression levels of *Orco* and *Ir25a* were normalized to the geometric mean of three reference genes (*gapdh*, *ppil2*, and *rpL13a*). The sequences of all qPCR primers are provided in table S5.

#### HCR RNA-FISH

##### OCT Embedding

Bees used in this experiment were derived from another SDI colony. Second instar larvae were removed from their colonies as described above and quickly rinsed with 1 x Dulbecco’s phosphate-buffered saline (DPBS). We then placed each larva flat on a small amount of OCT (Fisher HealthCare, USA) in a cryomold and then filled the mold completely with OCT. The cryomold was placed on a petri dish floating in liquid nitrogen to allow rapid bottom-up freezing until it turned opaque white. Finally, the cryomold was transferred to a −80°C freezer for subsequent processing.

##### Sectioning, Fixation, and Permeabilization

Second instar larvae were sectioned into 10 μm thick slices with a Leica cryostat (Leica CM3050S, Germany) and placed on 18 mm coverslips. The sections were dried for 1 min per micron of thickness, followed by fixation with 4% paraformaldehyde (PFA) in 1x phosphate-buffered saline (PBS) for 13 min. Samples were then washed three times with 1x PBS to remove the residual PFA. The tissues were then rinsed with cold 70% ethanol and permeabilized by incubating in 70% ethanol overnight at 4°C.

##### Probe Hybridization

Probes for *Orco*, *Ir25a*, and *Elav* (a neural-specific gene to provide general spatial information) were used for hybridization (Custom probes from Molecular Instruments). The probe hybridization buffer (Molecular Instruments, USA) was pre-warmed to 37° C before use. A humidified chamber, glass plates, and Parafilm® M were UV-treated for 60 min. The ethanol was then removed from the samples, followed by two 5-min washes with PBS-Tw (0.1% Tween-20 in 1× PBS) at room temperature. Before probe hybridization, the samples were incubated in the warm probe hybridization buffer for 10 min at 37°C. Next, the initiator probe solution was prepared by adding each stock probe to the warm probe hybridization buffer to a final concentration of 2 pmol per probe. Each tissue section was placed on 25 μL of the prepared initiator probe solution in a humidified chamber and incubated for 36-48 h at 37°C.

##### Amplification and Hybridization

Probe wash buffer was prewarmed to 37 ° C and amplification buffer to room temperature. The samples were washed with probe wash buffer (Molecular Instruments, USA), followed by a 1-h incubation at 37°C. The samples were then incubated in 5x SSCTw (0.1% Tween-20 in 5x SSC) buffer for 30 min at room temperature. Hairpin probes (h1 and h2) were prepared by adding 7 μL of 3 μM hairpin stock to each PCR tube. The tubes were incubated at 95°C for 90 sec and subsequently cooled in the dark at room temperature for at least 30 min. Then the amplifier hybridization buffer (Molecular Instruments, USA) was prepared by adding hairpins to the amplification buffer with a final concentration of 180 nM per hairpin. Each tissue section was then placed on 25 μL of amplifier hybridization buffer and incubated overnight in a humidified chamber at 37°C.

##### Wash and Staining

The next day, samples were washed with 5x SSCT (0.1% TritonX-100 in 5x SSC) buffer to remove excess amplification buffer and hairpins. Nuclei were stained with 10 μg/mL 4′,6-diamidino-2-phenylindole (DAPI) in 5x SSCT buffer for 10 minutes, followed by a final wash in 5x SSCT buffer for 5 minutes. The samples were then mounted onto glass plates for downstream imaging.

##### Imaging

Imaging was performed using a Nikon (Japan) Ti2-E widefield microscope equipped with four color channels: 405 nm for DAPI, 647 nm for *Elav*, 565 nm for *Orco*, and 465 nm for *Ir25a*. Exposure times for each channel were set to 50*μ*s, 300-500*μ*s, 100-300*μ*s, and 300*μ*s, respectively, depending on the signal intensity and dynamic range of each channel. The laser source was a Lumencor CELESTA light engine. A 5-band dichroic filter (Chroma) was used for all images. An sCMOS camera (Kinetix) was employed to acquire the images. Image acquisition and stitching were conducted using NIS-Elements. Samples were imaged with a 63x oil-immersion objective. Five z-planes were captured at 1.2 µm step sizes for each field of view. Post-acquisition analysis was also performed using the NIS software suite to ensure accurate data interpretation.

#### Comparative analyses of larval gene expression in honey bees

We compared the expression of *Orco* and *Ir25a* with data from seven published RNAseq studies of honey bee larvae (n = 21) (*16-22*), obtained from the SRA database using the fastq-dump tool from the SRA Toolkit (https://ftp-trace.ncbi.nlm.nih.gov/sra/sdk/2.10.7/). Sequencing quality was checked with FastQC v.0.11.8 (*46*). Data were aligned to the latest honey bee reference genome assembly HvA3.1 (*47*) by Salmon v1.4.0 (*48*) with default parameters. To evaluate the expression levels of each gene within each sample, we further computed the FPKM (fragments per kilobase of transcript per million mapped reads) values based on the length of the gene and the count of reads mapped to this gene (*49*).

#### Comparative analyses of larval gene expression in different insect species

We compared the expression of *Orco* and *Ir25a* with data from published RNAseq studies of larvae from 11 species: *A. mellifera* (*16-22, 27*); *Bombus terrestris* (*28*); *Monomorium pharaonis* and *D. melanogaster* (*27, 29*); *Megalopta genalis* (*30*); *Ceratina calcarata* (*31*); *Bactrocera dorsalis* (*32*); *Plutella xylostella* (*33*); *Zeugodacus cucurbitae*; *Bactrocera latifrons* (*34*); and *Anopheles gambiae* (*35, 36*). We quantified gene expression using FPKM.

#### Statistical analyses

For the behavioral analysis, which measured the total and cumulative number of food droplets consumed by each larva per hour over the course of each 17-hour experiment, we performed a Wilcoxon rank sum test. Spearman’s rank correlation tests were used to examine the relationship between larval weight and number of food droplets consumed.

For the qPCR analysis, differences in expression between developmental stages and colonies for all genes except *Rcp1* were analyzed using linear mixed-effects models (LME) with the nlme package 3.1-157 in the R environment version 4.2.1. This approach allowed for the incorporation of colony as a random effect to account for variability among colonies, with developmental stages treated as fixed effects. If necessary, data were log- or square root transformed to achieve a Gaussian distribution of the model residuals. After fitting the model, the significance of differences in expression levels across developmental stages was assessed. Then Tukey’s post hoc test was used for pairwise comparisons between developmental stages to identify specific differences.

**Fig. S1.**
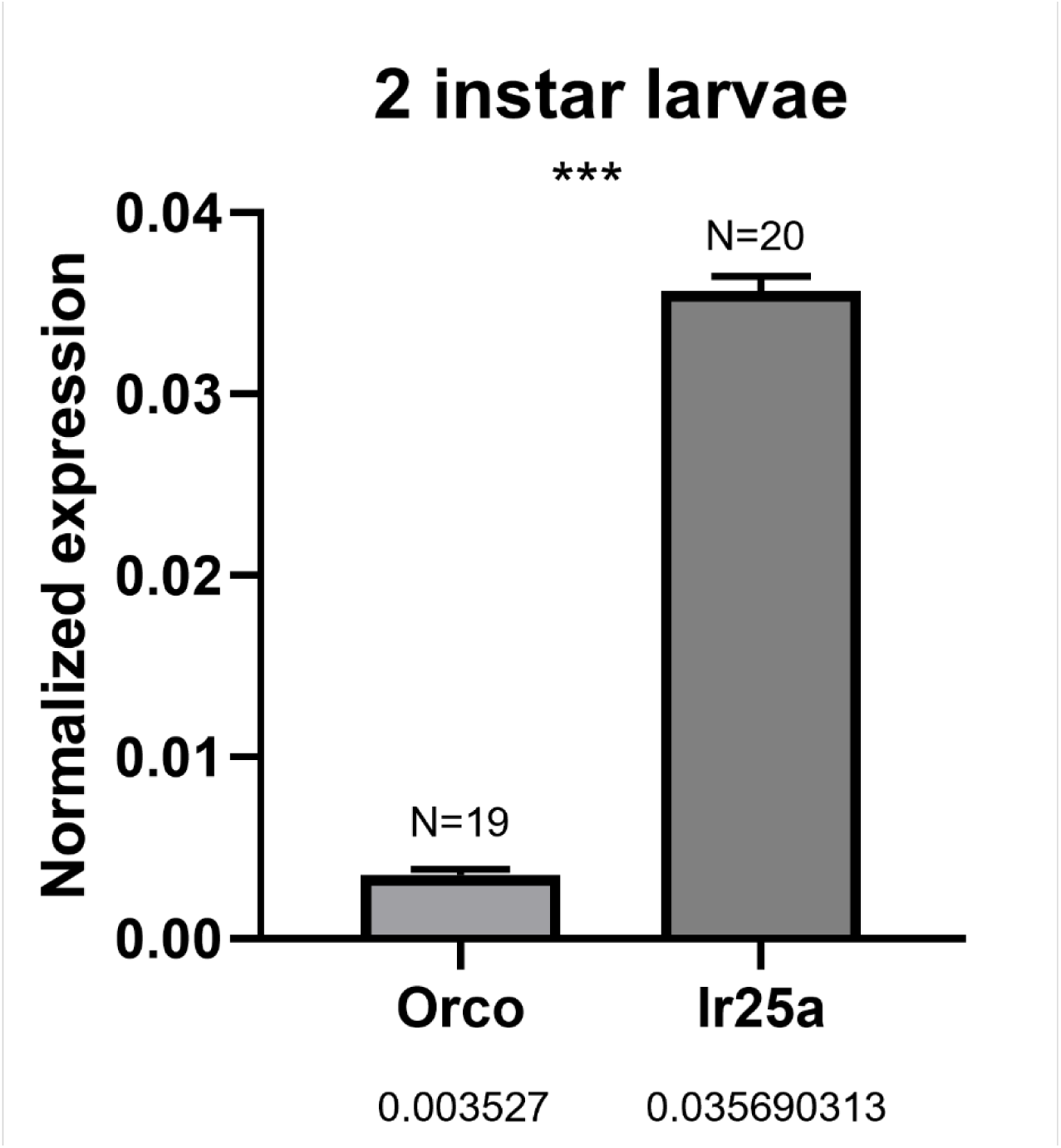
Data are presented as mean ± s.e. Numbers below the bars are the normalized expression values of each gene relative to a pool of 3 housekeeping genes. *** indicates significant differences between the expression levels of *Orco* and *Ir25a* (p < 0.0001).

**Fig. S2.**
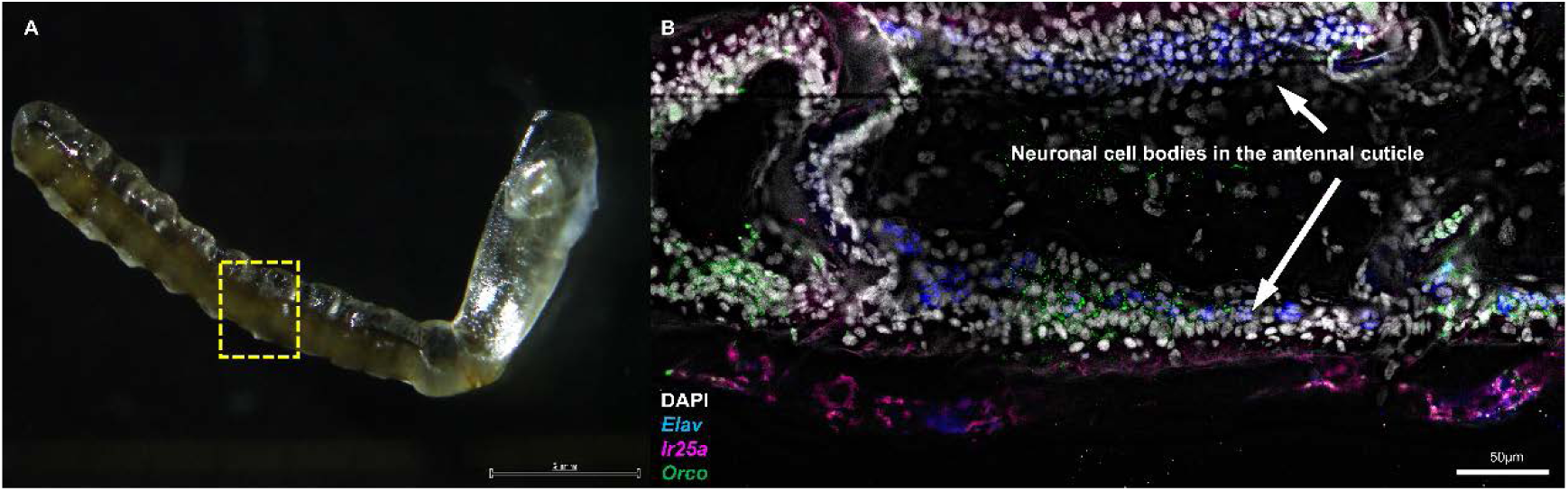
Expression of *Orco* and *Ir25a* in pupal antenna. HCR RNA-FISH was performed on longitudinal tissue sections of worker pupal antenna. (A) Lateral view of the pupal antenna. (B) Higher magnification of the area boxed in A shows hybridization in the longitudinal section of the pupal antenna. Different colors represent different genes as per legend. *Elav* was used as a neuronal marker, and DAPI was used as a stain for nuclear quantitation. Arrows indicate the region of neuronal cell bodies in the pupal antennal cuticle. Signals for both *Orco* and *Ir25a* were detected above the background level.

**Video files may be downloaded or viewed at the following links.**

**Video S1.** The trajectory of bee larvae moving in Petri dishes for 17 hours. https://drive.google.com/file/d/1VqEagUvp4xnZy3yUymdM9aEdw4ribrRQ/view?usp=drive_link

**Video S2**. The larva did not show any aversion to diet C and actively fed on it. https://drive.google.com/file/d/1-xkC_Rb5Pjn65d1CpWA3GYWQDpsksqMq/view?usp=sharing

## References

1. D. E. Vázquez, W. M. Farina, Locomotion and searching behaviour in the honey bee larva depend on nursing interaction. Apidologie 52, 1368–1386 (2021).

2. G. E. Robinson, Chemical communication in honeybees. Science 271, 1824–1825 (1996).

3. E. Oertel, Metamorphosis in the honeybee. Journal of Morphology 50, 295–339 (1930).

4. H. M. Robertson, K. W. Wanner, The chemoreceptor superfamily in the honey bee, *Apis mellifera*: expansion of the odorant, but not gustatory, receptor family. Genome Research 16, 1395–1403 (2006).

5. N. M. Tian, D. J. Price, Why cavefish are blind. Bioessays 27, 235–238 (2005).

6. J. J. Elser, C. Acquisti, S. Kumar, Stoichiogenomics: the evolutionary ecology of macromolecular elemental composition. Trends in Ecology & Evolution 26, 38–44 (2011).

7. J. Isanta-Navarro et al., Revisiting the growth rate hypothesis: Towards a holistic stoichiometric understanding of growth. Ecology Letters 25, 2324–2339 (2022).

8. G. Dhar et al., "Various behavioural assays to detect the neuronal abnormality in flies" in Fundamental approaches to screen abnormalities in Drosophila (Springer, 2020), pp. 223-251.

9. V. G. Dethier, Gustation and olfaction in lepidopterous larvae. The Biological Bulletin 72, 7–23 (1937).

10. C. C. Arce et al., Plant-associated CO_2_ mediates long-distance host location and foraging behaviour of a root herbivore. Elife 10, e65575 (2021).

11. V. Croset et al., Ancient protostome origin of chemosensory ionotropic glutamate receptors and the evolution of insect taste and olfaction. PLoS Genetics 6, e1001064 (2010).

12. L. Abuin et al., Functional architecture of olfactory ionotropic glutamate receptors. Neuron 69, 44–60 (2011).

13. H. Yan et al., An engineered *orco* mutation produces aberrant social behavior and defective neural development in ants. Cell 170, 736–747 (2017).

14. W. Trible et al., *orco* mutagenesis causes loss of antennal lobe glomeruli and impaired social behavior in ants. Cell 170, 727–735 (2017).

15. Z. Chen et al., Neurodevelopmental and transcriptomic effects of CRISPR/Cas9-induced somatic *orco* mutation in honey bees. Journal of Neurogenetics 35, 320–332 (2021).

16. D. E. Vazquez, J. M. Latorre-Estivalis, S. Ons, W. M. Farina, Chronic exposure to glyphosate induces transcriptional changes in honey bee larva: A toxicogenomic study. Environmental Pollution 261, 114148 (2020).

17. V. Kowallik, A. S. Mikheyev, Honey bee larval and adult microbiome life stages are effectively decoupled with vertical transmission overcoming early life perturbations. mBio 12, e0296621 (2021).

18. J. Luo et al., The comparison of juvenile hormone and transcriptional changes between three different juvenile hormone analogs insecticides on honey bee worker larval’s development. Agronomy 11, 2497 (2021).

19. H. Li et al., Juvenile hormone and transcriptional changes in honey bee worker larvae when exposed to sublethal concentrations of thiamethoxam. Ecotoxicology and Environmental Safety 225, 112744 (2021).

20. X. J. He, W. J. Jiang, M. Zhou, A. B. Barron, Z. J. Zeng, A comparison of honeybee (*Apis mellifera*) queen, worker and drone larvae by RNA-Seq. Insect Science 26, 499–509 (2019).

21. X. J. He et al., Starving honey bee (*Apis mellifera*) larvae signal pheromonally to worker bees. Scientific Reports 6, 22359 (2016).

22. X. J. He et al., Extent and complexity of RNA processing in honey bee queen and worker caste development. iScience 25, 104301 (2022).

23. H. Nie et al., Comparative transcriptome analysis of *Apis mellifera* antennae of workers performing different tasks. Molecular Genetics and Genomics 293, 237–248 (2018).

24. S. Stewart, T. W. Koh, A. C. Ghosh, J. R. Carlson, Candidate ionotropic taste receptors in the *Drosophila* larva. Proceedings of the National Academy of Sciences 112, 4195–4201 (2015).

25. H. Yan et al., Evolution, developmental expression and function of odorant receptors in insects. Journal of Experimental Biology 223, (suppl. 1), jeb208215 (2020).

26. E. Jongepier et al., Convergent loss of chemoreceptors across independent origins of slave-making in ants. Molecular Biology and Evolution 39, msab305 (2022).

27. M. R. Warner, L. Qiu, M. J. Holmes, A. S. Mikheyev, T. A. Linksvayer, Convergent eusocial evolution is based on a shared reproductive groundplan plus lineage-specific plastic genes. Nature Communications 10, 2651 (2019).

28. T. J. Colgan et al., Genomic signatures of recent adaptation in a wild bumblebee. Molecular Biology and Evolution 39, msab366 (2022).

29. S. Hu Qian et al., Integrating massive RNA-seq data to elucidate transcriptome dynamics in *Drosophila melanogaster*. Briefings in Bioinformatics 24, bbad177 (2023).

30. K. M. Kapheim et al., Developmental plasticity shapes social traits and selection in a facultatively eusocial bee. Proceedings of the National Academy of Sciences 117, 13615–13625 (2020).

31. K. D. Chau, M. Shamekh, J. Huisken, S. M. Rehan, The effects of maternal care on the developmental transcriptome and metatranscriptome of a wild bee. Communications Biology 6, 904 (2023).

32. E. H. Chen et al., RNA-seq analysis of gene expression changes during pupariation in *Bactrocera dorsalis* (Hendel) (Diptera: Tephritidae). BMC Genomics 19, 1–16 (2018).

33. Q. L. Hou, E. H. Chen, RNA-seq analysis of gene expression changes in cuticles during the larval-pupal metamorphosis of *Plutella xylostella*. Comparative Biochemistry and Physiology Part D: Genomics and Proteomics 39, 100869 (2021).

34. C. Liang et al., TephritidBase: a genome visualization and gene expression database for tephritid flies. Arthropod-Plant Interactions 18, 379–388 (2024).

35. G. Rose et al., Dosage compensation in the African malaria mosquito *Anopheles gambiae*. Genome Biology and Evolution 8, 411–425 (2016).

36. C. M. Mang’era et al., Transcriptomic response of *Anopheles gambiae* sensu stricto mosquito larvae to Curry tree (*Murraya koenigii*) phytochemicals. Parasites & Vectors 14, 1–14 (2021).

37. M. C. Larsson et al., *Or83b* encodes a broadly expressed odorant receptor essential for *Drosophila* olfaction. Neuron 43, 703–714 (2004).

38. J. E. Niven, S. B. Laughlin, Energy limitation as a selective pressure on the evolution of sensory systems. Journal of Experimental Biology 211, 1792–1804 (2008).

39. C. Gadenne, R. B. Barrozo, S. Anton, Plasticity in insect olfaction: to smell or not to smell? Annual Review of Entomology 61, 317–333 (2016).

40. T. Giray, G. E. Robinson, Effects of intracolony variability in behavioral development on plasticity of division of labor in honey bee colonies. Behavioral Ecology and Sociobiology 35, 13–20 (1994).

41. C. W. Whitfield, A.-M. Cziko, G. E. Robinson, Gene expression profiles in the brain predict behavior in individual honey bees. Science 302, 296–299 (2003).

42. F. Wu et al., Behavioural, physiological and molecular changes in alloparental caregivers may be responsible for selection response for female reproductive investment in honey bees. Molecular Ecology 28, 4212–4227 (2019).

43. D. Task et al., Chemoreceptor co-expression in *Drosophila melanogaster* olfactory neurons. Elife 11, e72599 (2022).

44. H. Yan, Insect olfactory neurons: receptors, development, and function. Current Opinion in Insect Science 67, 101288 (2025).

45. I. M. Traniello et al., Context-dependent influence of threat on honey bee social network dynamics and brain gene expression. Journal of Experimental Biology 225, jeb243738 (2022).

46. S. Anders, P. T. Pyl, W. Huber, HTSeq--a Python framework to work with high-throughput sequencing data. Bioinformatics 31, 166–169 (2015).

47. K. L. Howe et al., Ensembl Genomes 2020-enabling non-vertebrate genomic research. Nucleic Acids Research 48, D689–D695 (2020).

48. R. Patro, G. Duggal, M. I. Love, R. A. Irizarry, C. Kingsford, Salmon provides fast and bias-aware quantification of transcript expression. Nature Methods 14, 417–419 (2017).

49. A. Conesa et al., A survey of best practices for RNA-seq data analysis. Genome Biology 17, 1–19 (2016).

